# Entering the era of conservation genomics: Cost-effective assembly of the African wild dog genome using linked long reads

**DOI:** 10.1101/195180

**Authors:** Ellie E. Armstrong, Ryan W. Taylor, Stefan Prost, Peter Blinston, Esther van der Meer, Hillary Madzikanda, Olivia Mufute, Roseline Mandisodza, John Stuelpnagel, Claudio Sillero-Zubiri, Dmitri Petrov

## Abstract

A high-quality reference genome assembly is a valuable tool for the study of non-model organisms across disciplines. Genomic techniques can provide important insights about past population sizes, local adaptation, and even aid in the development of breeding management plans. This information can be particularly important for fields like conservation genetics, where endangered species require critical and immediate attention. However, funding for genomic-based methods can be sparse for conservation projects, as costs for general species management can consume budgets. Here we report the generation of high-quality reference genomes for the African wild dog (*Lycaon pictus*) at a low cost, thereby facilitating future studies of this endangered canid. We generated assemblies for three individuals from whole blood samples using the linked-read 10x Genomics Chromium system. The most continuous assembly had a scaffold N50 of 21 Mb, a contig N50 of 83 Kb, and completely reconstructed 95% of conserved mammalian genes as reported by BUSCO v2, indicating a high assembly quality. Thus, we show that 10x Genomics Chromium data can be used to effectively generate high-quality genomes of mammal species from Illumina short-read data of intermediate coverage (∼25-50x). Interestingly, the African wild dog shows a much higher heterozygosity than other species of conservation concern, possibly as a result of its behavioral ecology. The availability of reference genomes for non-model organisms will facilitate better genetic monitoring of threatened species such as the African wild dog. At the same time, they can help researchers and conservationists to better understand the ecology and adaptability of those species in a changing environment.

## Introduction

Major population declines have been observed in vertebrate groups over the past several hundred years, primarily due to anthropogenic change (Pimm et al. 2014). This decline has resulted in extinction rates unprecedented in recent history (Pimm et al. 2014; Ceballos et al. 2015). The conservation of extant species will require major efforts in restoring and preserving habitat, along with protection, management, and investment by local stakeholders. Though many species of conservation concern exist as small populations, populations can still retain genetic variation that was generated and maintained a few generations back, when population sizes were much larger. Within patterns of historic genetic variation are signals of demographic history, gene flow, and natural selection which can inform efforts towards the long-term survival of species. In addition to signals of a species history, genetic information can be used to uncover important contemporary or very recent events and processes. For example, Epstein et al. (2016) identified genes that may confer facial tumor resistance in Tasmanian devils, suggesting that the ability to artificially select for resistance in non-infected populations may allow for a more robust population rescue and recovery. Genetic markers can be used to track individual movement across landscapes either indirectly by measuring relatedness, or directly by genotyping scat or hair left by an individual as it moves. Additionally, the identification and assignment of individuals through genotyping can be an important tool for law enforcement to assign contraband and confiscated materials to their geographic origin. Conservationists can also use fine grained measurements of reproductive success along with genotypes and environmental variables to gather a detailed understanding of the factors contributing to or limiting population growth, such as inbreeding depression. Taken together genomic tools are poised to have a major contribution to conservation (Steiner et al. 2013; Shafer et al. 2015).

The African wild dog (*Lycaon pictus*) is a medium-sized (18-34kg), endangered carnivore that lives in scattered populations in sub Saharan Africa (Fig. 1A). The species is the only surviving member of a lineage of wolf-like canids (Girman et al. 1993). Wild dogs have been subject to intense recovery efforts across its range (Woodroffe et al. 1997; IUCN/SSC 2007), but their global population is decreasing. It is estimated that only 6,600 adult wild dogs remain in 39 subpopulations (Woodroffe & Sillero-Zubiri 2012). The primary reasons for the species’ population decline include habitat loss and fragmentation, as well as anthropogenic mortality (e.g. snaring, persecution, road kills, exposure to infectious diseases from domestic dogs) when they range beyond the borders of protected areas (Woodroffe et al. 1997; Woodroffe & Ginsberg 1998; IUCN/SSC 2007). Due to their large ranges and low population densities, African wild dogs are more susceptible to these threats than most other carnivore species (IUCN/SSC 2007). In addition, their complex social system and susceptibility to Allee effects appears to increase the species extinction risk (Courchamp et al. 1999, 2000). The dogs are obligate cooperative breeders which form packs consisting of an alpha male and female, their adult siblings, and pups and subadults from the dominant pair (McNutt & Silk 2008). Subadults that have reached reproductive age disperse in single sex groups and form new packs by joining dispersing groups from the opposite sex (McNutt 1996). Pack members rely on each other for hunting, breeding, and defense against natural enemies and pack size has been found to be significant for hunting and breeding success (Fanshawe & Fitzgibbon 1993; Creel & Creel 1998; McNutt & Silk 2008). When pack size becomes critically low, e.g. due to anthropogenic mortality, this dependence on helpers increases the risk of pack extinction and reduces the number of successful dispersals (Courchamp et al. (1999) and Courchamp et al. (2000), but see Creel and Creel (2015)).

**Figure 1.**
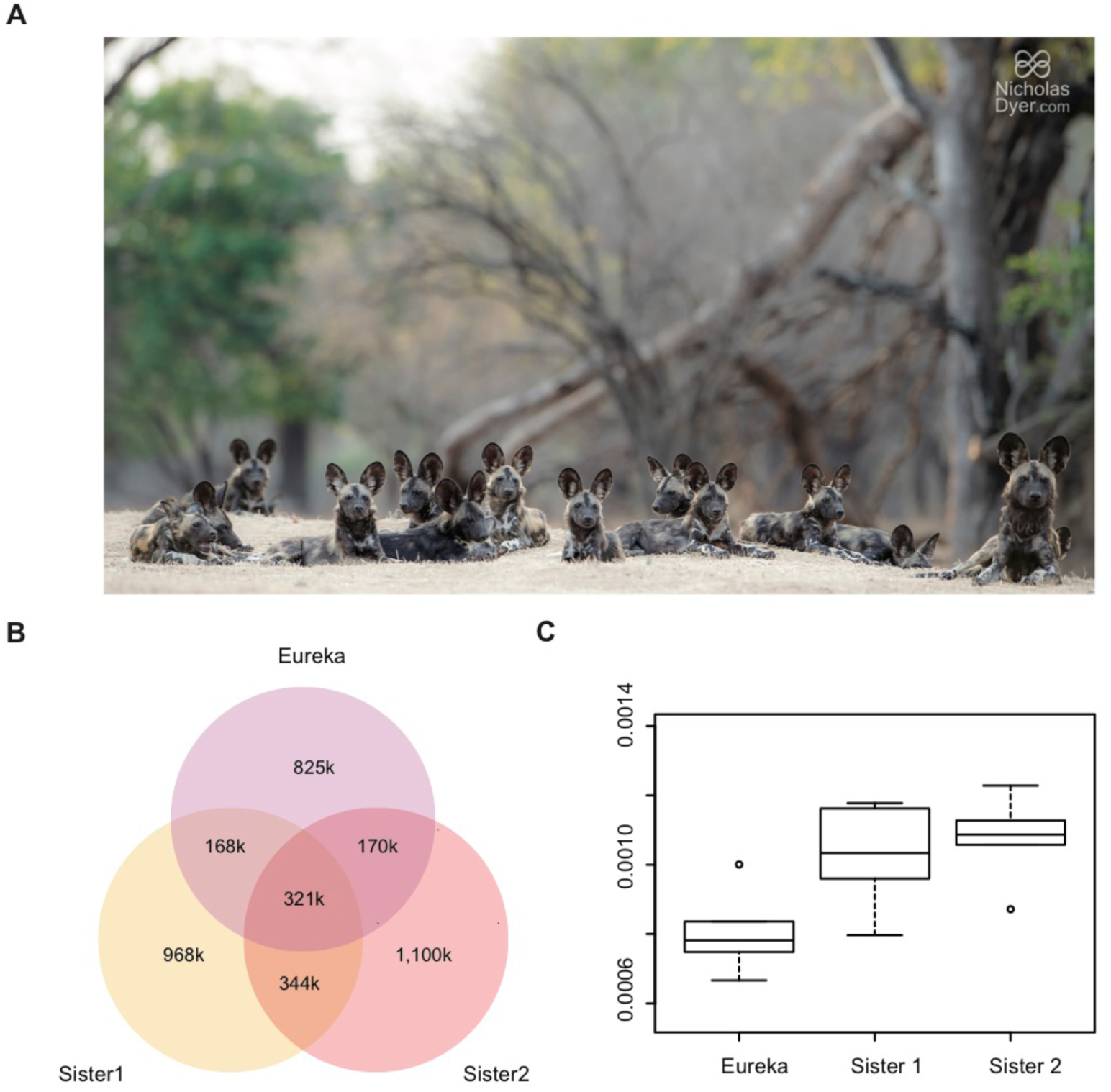
Shared heterozygous sites between the different African wild dog individuals. A) Pack of African wild dogs. B) Shared heterozygous sites between the three *de novo* assemblies (calculated using a posterior cutoff of 0.99). Many of the heterozygous sites are shared between all individuals and more heterozygous sites are shared between the two sisters than between each sister and Eureka. C) Boxplot of heterozygosity values (y-axis) calculated for different posterior probability cutoffs.

Prior genetic studies on wild dogs using a combination of mitochondrial, microsatellite, and MHC markers have resulted in varying estimates of the start of the species decline on the African continent (Girman et al. 2001; Marsden et al. 2012). Consistent with expectation, the data shows strong structuring between populations due to habitat fragmentation and isolation, as well as low genetic diversity within populations (Marsden et al. 2009; Marsden et al. 2012). For species that are experiencing such rapid and alarming declines, estimates that are particularly important for management decisions, such as local adaptation, effective population size, and inbreeding are greatly improved by the use of whole-genome methods. Recently, Campana and colleagues (Campana et al. 2016) sequenced low-coverage genomes of two African wild dog individuals from Kenya and South Africa, respectively, to investigate demographic history and signatures of selection of these two separate populations. By mapping these data to the domestic dog genome, they discovered approximately 780,000 single nucleotide polymorphisms (SNPs) between their two individuals which could be used to develop SNP typing for the two populations. However, given the low coverage of their genomes (5.7-5.8x average coverage) and the small number of individuals, additional sequencing will be needed to verify the authenticity of those SNPs. Further, important structural variation can be overlooked when mapping against a reference genome from a different genus, and mapping can be hindered if the divergence is high between the sample and the reference (see e.g. Lunter and Goodson (2011), Shapiro and Hofreiter (2014)). The groups containing the African wild dog and the domestic dog are estimated to have split approximately 7.5-10 Mya and furthermore, the domestic dog has undergone significant genomic selection in recent time (Nyakatura & Bininda-Emonds 2012).

Despite the ever-declining cost to sequence DNA, the routine use of genomic approaches in conservation is still far from a reality. One of the major remaining barriers is the lack of reference genomes for species of conservation concern. Generating a *de novo* reference genome requires the sequencing and assembly of the 100s of millions to billions of base-pairs that make up a genome. The first mammalian genome (human) required a massive collaboration between hundreds of scientists and nearly $3 billion US dollars (1990-2001; (Lander et al. 2001; Hayden 2014)). Fortunately, the cost to sequence DNA is now low enough that every base-pair in a typical mammalian genome can be sequenced to high coverage for a few thousand US dollars. However, these low cost sequencing methods produce very short sequences of 150-300 base-pairs in length (for a review on sequencing methods see Goodwin et al. (2016)). Because large proportions of typical mammal genomes consist of repetitive sequences, it has been impossible to assemble highly-contiguous genomes from only these short sequences. In order to achieve higher continuity, more elaborate and expensive library preparation or alternative sequencing technologies have to be used (Ekblom & Wolf 2014; Goodwin et al. 2016). Among others, these include mate-pair libraries, chromatin folding based libraries, such as cHiCago (Putnam et al. 2016) or HiC (Burton et al. 2013), and long-read sequencing technologies, such as Pacific Biosciences and Oxford Nanopore Technology. While the resulting genomes can show high continuity, those methods substantially increase the costs of sequencing projects and thus can hinder the generation of genomes for conservation biology purposes.

Here we report the use of the Chromium system developed by 10x Genomics (Weisenfeld et al. 2017), a genomic library preparation technique that facilitates cost-effective (around $2,500) assemblies using short sequencing reads, to assemble three African wild dog genomes. In brief, the 10x Genomics Chromium system is based on dilution of high molecular weight (HMW) DNA. It uses as little as 1ng of input DNA, which is well-suited for a variety of applications. During library preparation, gel beads, so-called GEMs, are mixed with DNA and polymerase for whole-genome amplification. Each gel bead has primer oligos (44nt long) attached to its surface. These contain a priming site (22nt partial R1), a 16nt barcode region, and a 6nt N-mer region that binds to different places on the original DNA fragment. The low amount of input DNA ensures that each gel bead only binds a single (up to ∼100kb) DNA fragment. In the next step, amplification of short reads along the original DNA fragment is performed within each gel bead. In most cases, this amplification results in spotted read coverage along the fragment. However, all reads from a respective GEM contain identical barcodes and can later be assigned to groups originating from the same DNA molecule. The information about which molecule of DNA the sequence originated from greatly increases the ability to identify the location of repetitive sequences. The library is then sequenced on an Illumina platform and the raw read data is assembled by the 10x Genomics Supernova assembler. This assembler is very user-friendly and does not require any prior knowledge about input parameters for the assembly.

We *de novo* assembled three African wild dog genomes using the 10x Genomics Chromium platform in order to investigate whether this technology is suitable for conservation genomic purposes. For any endangered species, a genome can have large conservation impacts, but high-quality genomes have historically been costly or impossible due to the sampling requirements and in addition, downstream analyses can be challenging. Thus, in order for it to be useful for conservation purposes the technology needs to be (a) cost-effective and (b) user-friendly. Furthermore, we test the 10x Genomics Chromium based assemblies for reproducibility, continuity, conserved gene completeness, and repetitive content, as compared to the previously published domestic dog genome.

## Methods

### Samples

Blood samples from two individuals belonging to the same pack in Hwange National Park, Zimbabwe were provided by Painted Dog Conservation. These individuals were presumed to be sisters from direct observation of their litter at the den (here, named Sister 1 and Sister 2). Both samples were collected during routine collaring and health monitoring. From these samples 3ml of blood was aliquoted and frozen immediately in liquid nitrogen and kept frozen in liquid nitrogen for 6 months until transfer to a -80°C freezer. DNA was extracted two weeks after storage at - 80°C. The third sample was provided by the Endangered Wolf Center, Eureka, Missouri from a captive born individual (here named Eureka). This individual’s mother descended from a male that was wild caught in Ellisras/Lephalale in the Limpopo Province of South Africa, and from a female that was wild caught in Botswana (no further details about location are available). The blood from this sample was treated with EDTA anticoagulant, refrigerated, and shipped on ice. DNA was extracted 9 days after the sample was taken. Though the Chromium library preparation does not require large amounts of DNA, the DNA should have a mean molecule length > 200kb (high-molecular weight, or HMW). DNA from all individuals was extracted from blood samples using the QIAGEN MagAttract HMW DNA kit following the provided instructions.

### Genome Assembly

We constructed one sequencing library per individual using the 10x Genomics Chromium System with 1.2ng of HMW input DNA. The libraries for Sister 1 and Eureka were prepared and sequenced by 10x Genomics in Pleasanton, California. The library for Sister 2 was prepared and sequenced by HudsonAlpha in Huntsville, Alabama. All libraries were then sequenced on the Illumina HiSeqX (Sister 2, Eureka) or HiSeq 4000 (Sister 1) platform. We generated 1,200 million read pairs for Sister 1, 801.56 million reads for Sister 2, and 427.6 million reads for Eureka.

We subsequently assembled the three genomes using the 10x Genomics genome assembler Supernova 1.1.1 Weisenfeld et al. (2017); http://support.10xgenomics.com/de-novo-assembly/software/overview/welcome) using default assembly parameters.

### Assembly Quality Assessment

We used the Supernova assembler as well as QUASTv4.3 to determine continuity statistics, such as the scaffold N50 and the total number of scaffolds (Gurevich et al. 2013). In order to estimate the N50 statistic, scaffolds first get ranked according to their size. The N50 value is the size of the scaffold when the running sum (starting with the longest scaffold) equals at least half the genome size. It is similar to the median scaffold length, but puts more weight on longer scaffolds. We further applied the program BUSCO v2 (Simão et al. 2015) to assess the presence of nearly universal lineage specific single-copy orthologous genes in our assemblies using the mammalian gene set from OrthoDB v9 (4104 genes; available at http://busco.ezlab.org). We compare these results to the high-quality canFam3.1 assembly of the domestic dog (Hoeppner et al. (2014); *Canis familiaris*). The canFam3.1 assembly was built on 7x coverage of Sanger reads and BAC end sequencing and has a scaffold N50 of 46Mb. Prior to long-read technology, this approach was the gold standard to generate high-quality genomes of model organisms. This approach is especially useful for resolving repetitive or complex regions, but unfortunately it is very costly. We also estimated the number of BUSCO’s using the recently published Hawaiian monk seal genome (which was assembled using a combination of 10x Genomics Chromium and Bionano Genomics Irys data and the two previously published African wild dog genomes (sequenced with basic short read Illumina technology at low coverage and assembled using the domestic dog; (Campana et al. 2016)).

### Repeat Identification and Masking

We next identified repetitive regions in the genomes as another comparative measure of assembly quality and to prepare the genome for annotation. Repeat annotation was carried out using both homology-based and *ab-initio* prediction approaches. We used the canid RepBase (http://www.girinst.org/repbase/; (Jurka et al. 2005)) repeat database for the homology-based annotation within RepeatMasker (http://www.repeatmasker.org; (Smit et al.)). In this step, previously compiled repeats from the canid database were mapped to the genome to identify repeats in the sequence. The RepeatMasker option -gccalc was used to infer GC content for each contig separately to improve the repeat annotation. We then carried out *ab-initio* repeat finding using RepeatModeler (http://repeatmasker.org/RepeatModeler.html; (Smit et al. 2014)). On the contrary to RepeatMasker, RepeatModeler does not require previously assembled repeat databases, but identifies repeats in the genome using statistical models.

### Gene Annotation

Gene annotation for the three assemblies was performed with the genome annotation pipeline Maker3 (Holt & Yandell 2011), which implements both *ab-initio* prediction and homology-based gene annotation by leveraging previously published protein sequences from dog, mouse, and human. In order to reduce the number of false positives, we hard-masked tandem elements before running the pipeline. Hard-masking replaces repeat sequences with Ns and thereby precludes any alignment to these regions. On the other hand, we only soft-masked simple repeats (conversion of sequences to lowercase). This allows alignment to these regions, but prevents the simple repeat from being included in the gene model during the actual gene annotation. We configured Maker3 to soft-mask simple repeats during the pipeline run.

Orthologous genes between the three African wild dog assemblies, as well as paralogous genes within each individual, were inferred using proteinortho (Lechner et al. 2014). Proteinortho applies highly parallelized reciprocal blast searches to establish orthology and paralogy for genes within and between gene annotation files.

### Variant rates

In order to estimate within individual heterozygosity, we selected a single pseudo-haplotype (in cases where genomic regions were phased into haplotypes, one of the two was chosen randomly) from Sister 2 to represent the reference sequence. Next we mapped the raw reads from all three individuals to the reference using bwa mem (Li & Durbin 2009). We then converted the resulting sam files to bam format using samtools (Li et al. 2009), and sorted and indexed them using picard (http://broadinstitute.github.io/picard/). Realignment around insertion/deletion (indel) regions was performed using GATK, and finally, we called heterozygous sites using a probabilistic framework implemented in ANGSD (Korneliussen et al. 2014). We choose a probabilistic over a simple allele counting approach for two reasons. First, a genome coverage of 20x is on the lower side of what is needed to reliably call genotypes (Nielsen et al. 2012). However, even if coverage is as high as 55x, heterozygous sites can be falsely called due to erroneous alignment in low-complexity regions or if reads span areas not covered by the reference genome (Li 2014). Showing that even high coverage data could benefit from the application of probabilistic genotype calling. Here, we further addressed the former issue by applying realignment around indel regions using GATK. Second, we wanted to use the same approach for all samples, including the low coverage ones from Campana et al. (2016). We tested different posterior probability cutoffs (1, 0.999,0.99 and 0.95) using -doPost 2 -doCutoff 0.95 (with the following filters: -minIndDepth 15 - only_proper_pairs 0 -minQ 20). For the two genomes from Campana et al. (2016) we applied -minIndDepth 3 (given their average coverage of 5.7-5.8x). To allow for comparison between all individuals, we down-sampled all individuals to 20x mean nominal coverage (total number of reads covering a position, independent of their barcode) for our analyses. Heterozygosity was then simply calculated as the ratio of variable sites to the total number of sites (variable and invariable). Furthermore, Supernova outputs the distance between heterozygous sites as part of their assembly report. Briefly, here heterozygous sites are called from the assembly graph and are used for phasing (to generate a diploid genome consensus). We further downloaded the read data of Campana et al. (2016) and mapped them against our Sister 2 assembly to compare heterozygosity estimates (using the approach outlined above). We further estimated the number of shared heterozygous sites between our individuals. To do so, we used the *gplots* library in R (https://www.r-project.org) to calculate the overlap between the three sets and to display them in a Venn diagram. We then also integrated the two individuals from Campana et al. (2016) in this analysis. However, it is important to point out that those genotype calls are based on low-coverage data and may not be reliable (see e.g. (Nielsen et al. 2011; Nielsen et al. 2012)).

## Results

### Assembly of the African wild dog genome

Using 10x Genomics Chromium technology, we generated DNA libraries for three African wild dog individuals, two of which were collected from a wild pack in the Hwange National Park, Zimbabwe and are presumed to be sisters (named Sister 1 and Sister 2), and a third unrelated individual from the Endangered Wolf Center, Eureka, Missouri (named Eureka). A summary of the assembly statistics output by the Supernova assembler can be found in Table 1 (detailed statistics for each genome assembly can be found in Supplementary Table 1). We generated 1,200 million paired-end reads for Sister 1, 801.56 million reads for Sister 2, and 427.6 million reads for Eureka. We then used the reads to assemble each genome using the 10x Genomics Supernova assembler (as explained in https://support.10xgenomics.com/de-novo-assembly/software/overview/welcome). The mean input DNA molecule length reported by the Supernova assembler for Sister 1 was 19.91kb, Sister 2 was 77.03kb, and Eureka was 52.00kb. All three assemblies corroborate a genome size of approximately 2.3Gb, which is similar to that of the domestic dog (2.4Gb). These three assemblies together constitute the first reported *de novo* assemblies for the African wild dog species.

**Table 1.** Assembly Statistics. Assembly statistics for the three African wild dog genomes reported by the Supernova assembler. Coverage was assessed using samtools depth.

|  |  | **Sister 1** | **Sister 2** | **Eureka** |
| --- | --- | --- | --- | --- |
| **Input** | **Reads (m)** | 1,200 | 801.56 | 427.6 |
|  | **Average Coverage** | 69 | 46 | 25 |
|  | **Mean molecule size (kb)** | 19.91 | 77.03 | 52.00 |
| **Contig** | **N50 (kb)** | 61.34 | 83.47 | 50.15 |
|  | **Longest (kb)** | 524.60 | 615.40 | 450.50 |
|  | **Number (k)** | 78.62 | 68.64 | 108.00 |
| **Scaffold** | **N50 (mb)** | 7.91 | 21.34 | 15.31 |
|  | **Longest (kb)** | 43.96 | 69.63 | 41.67 |
|  | **Number (k)** | 11.78 | 17.64 | 25.78 |
| **Total Size (gb)** | **Scaffolds >= 10kb** | 2.27 | 2.26 | 2.20 |
|  | **Scaffolds >= 500bp** | 2.34 | 2.40 | 2.42 |

We then calculated the scaffold and contig N50 statistics, which are indicative of assembly continuity. The Sister 1 assembly resulted in a contig and scaffold N50 of 61.34 kb and 7.91 Mb, respectively, the Sister 2 assembly achieved 83.47 kb contig and 21.34 Mb scaffold N50s, and finally the Eureka assembly had 50.15 kb contig and 15.31 Mb scaffold N50s (Table 1). While our contig and scaffold N50’s are smaller than the ones from the most recent dog genome (267kb and 45.9Mb, respectively), they are still larger than most mammalian genomes assembled that used only short read data (see e.g. Figueiró et al. (2017) and Lok et al. (2017)).

### Conserved Genes

The program BUSCO (Benchmarking Universal Copy Orthologs) uses highly conserved single copy orthologous genes from a number of different taxa and groups in order to test assemblies (both genomic and transcriptomic) for gene completeness, fragmentation, or absence as an indicator of assembly quality. Using BUSCO v2 on our assemblies, we found that the most continuous assembly, Sister 2, completely recovered 95.1% of conserved genes (mammalia gene set; Table 2). Sister 1 and Eureka recovered 95.4% and 93.3% of complete conserved genes, respectively. Using the same analysis, we found 95.3% of complete conserved genes in the latest dog assembly (canFam3.1). This indicates that although the domestic dog assembly is more continuous overall, our assemblies recover nearly the same or even higher number of conserved genes. Surprisingly, Sister 1 had the least number of missing genes out of all the assemblies assessed, despite lower continuity than Sister 2. We also ran BUSCO on the Hawaiian monk seal genome, generated through the combination of 10x Genomics Chromium and Bionano Genomics Irys data, and found it recovered 94.6% of conserved genes using BUSCO. This suggests that using Bionano in addition to 10x does not greatly improve the ability reconstruct gene regions. However, the Hawaiian monk seal genome has a scaffold N50 of approximately 28Mb, so Bionano may improve the overall assembly continuity compared to 10x Genomics alone. The low coverage genomes from Campana et al. 2016 achieved a BUSCO score of 92.8% for the individual from Kenya and 94.8% for the individual from South Africa.

**Table 2.** Conserved Gene Statistics. Results of the BUSCO v2 gene annotation. The best values are shown in bold. We found that even though Sister 1 had lower continuity scores than Sister 2, this assembly recovered the most mammal orthologs (conserved genes). We also included the two individuals from Campana et al. (2016) and the Hawaiian Monk seal genome (Mohr et al. (2017); also assembled using the 10x chromium platform) for comparison.

| **Assembly** | **Species** | **Complete** | **Single Copy** | **Duplicated** | **Fragmented** | **Missing** | **Total Searched** |
| --- | --- | --- | --- | --- | --- | --- | --- |
| Sister 1 | *L. pictus* | **3914** | **3875** | 39 | 102 | **88** | 4104 |
| Sister 2 | *L. pictus* | 3903 | 3845 | 58 | 107 | 94 | 4104 |
| Eureka | *L. pictus* | 3829 | 3789 | 40 | 169 | 106 | 4104 |
| canFam3.1 | *C. familiaris* | 3910 | 3857 | 53 | **98** | 96 | 4104 |
| Kenya | *L. pictus* | 3849 | 3823 | 26 | 136 | 119 | 4104 |
| South Africa | *L. pictus* | 3892 | 3867 | **25** | 104 | 108 | 4104 |
| Hawaiian monk seal | *Neomonachus schauinslandi* | 3881 | 3833 | 48 | 118 | 105 | 4104 |

### Repeat annotation

We identified repetitive regions of the genome in order to discern how well these complex areas were assembled by the 10x Genomics Chromium technology. Using both RepeatMasker and RepeatModeler, we found that for all three wild dog assemblies, total repeat content was evaluated to be within 3% of one another, which indicates consistency among assemblies from a single species (Supplementary Table 2). No single repeat category was disproportionately affected during repeat annotation of the three genomes, which suggests that assembly quality was likely the most influential factor. Furthermore, repeat content of all wild dog assemblies was qualitatively similar to canFam3.1. As repetitive regions tend to be the most difficult regions to assemble, the similarity in repeat content between the wild dog compared to that of the domestic dog, highlights the value of using 10x Genomics Chromium technology to produce accurate and continuous assemblies.

### Gene annotation

The genome annotation pipeline Maker3 resulted in very similar numbers of annotated genes between all three wild dog individuals and the domestic dog. Annotations ranged from 20,649 (Sister 2) to 20,946 (Sister 1) genes (Supplementary Table 3). Using proteinortho to detect orthologous genes between individuals and paralogous genes within individuals, we found 12,617 one:one orthologs present in all three individuals and 6,462 one:one orthologs in two out of the three individuals. We found 268 multi copy genes present in all three individuals and 37 not present in one individual. Overall, the number of annotated genes was comparable to those found in the dog genome (Supplementary Table 3).

### Variant rates

We found a high number of heterozygous sites to be shared between all three individuals (321k; here we report the heterozygous sites called using a posterior probability cutoff of 0.99; Fig. 1B). As expected, Sister 1 and Sister 2 share more heterozygous sites (344k) than either sister with Eureka (168k and 170k, for Sister 1 and Sister 2, respectively). Each individual shows a high number of singletons (heterozygous sites only found in one individual), with Sister 2 showing the highest number (1,100k), followed by Sister 1 (968k) and Eureka (825k). Even if we include the two low coverage genomes from Campana et al. (2016), we find a high number of shared heterozygous sites between all individuals (134k; Supplementary Fig. 1). As expected, we see a higher number of singletons in these two individuals, due to the lower reliability of the genotype calls caused by the low coverage (false positives caused by sequencing errors). We estimated a per site heterozygosity of 0.0008 to 0.0012 for Sister 1, 0.0009 to 0.0012 for Sister 2, and 0.0007 to 0.001 for Eureka using posterior cutoffs for genotype calls from 0.95 to 1 in ANGSD (Supplementary Table 4; Fig. 1C). As can be seen in Supplementary Figure 2, except for a posterior probability cutoff of 1, where Sister 1 shows the highest heterozygosity, Sister 2 always shows the highest, Sister 1 the second highest and Eureka the lowest heterozygosity. Interestingly, Eureka shows a lower heterozygosity than the other two assemblies, even though its parents originated from South Africa and Botswana. Our estimates show that, while being heavily threatened, African Wild dogs can seem to still retain a relatively high within individual heterozygosity. We did not see any major difference between heterozygosity estimates from repeat-masked and unmasked genomes. The Supernova software estimated a heterozygous position every 2.6kb, 3.1kb, and 7.14kb for Sister 1, Sister2, and Eureka, respectively (Supplementary Table 1). On the contrary, estimates based on genotype calls using ANGSD showed much more frequent heterozygous positions (850bp - 1.2kb, 814bp - 1.1kb and 999bp - 1.5kb depending on the posterior cutoff used; Supplementary Table 4).

## Discussion

### Assembly continuity and quality

All three African wild dog assemblies produced with 10x Genomics Chromium data showed high continuity, high recovery rates of conserved genes, and expected proportions of repetitive sequence; indicating that they are high-quality assemblies. The Sister 2 assembly, which has the highest mean molecule length, is also the most continuous (Contig N50: 83.47kb, Scaffold N50: 21.34Mb; Table 1). Interestingly, the Sister 1 genome has a higher contig N50 (61.34kb) than Eureka (50.15kb), but a lower scaffold N50 (7.91Mb and 15.31Mb, respectively). This may indicate that input molecule length is a key factor for scaffolding, while coverage is a key factor for contig assembly. Despite having the highest continuity of all three assemblies, Sister 2 did not show the highest BUSCO completeness scores (see Table 2), although the differences were minor and likely not meaningful (with 95.1% complete BUSCOs compared to 95.4% for Sister 1). Sister 1 achieved the highest BUSCO scores, even compared to the latest domestic dog genome assembly (CanFam3.1; 95.2%), which has three times higher contig N50 and an almost six times higher scaffold N50. The high scores are remarkable for the limited number of reads used for the assemblies (as low as 25x coverage). As expected, Sister 2, which showed the highest continuity also had the highest repeat content (see Supplementary Table 2). However, all three assemblies resulted in similar repeat contents in terms of repeat composition as well as overall percentage (within 3% of each other), with the most continuous assembly (Sister 2) showing the highest number of repeats. Repeat composition in the African wild dog genomes was also similar to the domestic dog.

All assemblies yielded similar amounts of genes, with Sister 1 showing the highest number (see Supplementary Table 3), which reflects its BUSCO scores. Closer investigations of one:one and one:many orthologs further showed a very good agreement between annotations obtained from all three individuals. The numbers of annotated genes for all three African wild dogs were similar to those calculated for the latest domestic dog assembly.

### 10x Genomics Chromium system: Feasibility and caveats

Most mammal genomes published in the last several years use a mixture of paired-end (PE) and multiple mate pair (MP) Illumina libraries (e.g. Figueiró et al. (2017), Lok et al. (2017) and Liu et al. (2014)). While often resulting in good continuity (e.g. Liu et al. (2014) or Huang et al. (2014)), using different insert libraries considerably increases the cost per genome. On the contrary, 10x Genomics Chromium allows for assembly of a comparable or even more continuous genome using only a single library for a fraction of the cost (see below). Furthermore, as we show here, this library technology generates high-quality assemblies from as low as 25x coverage (see Eureka assembly), while the recommended coverage for PE plus MP assemblies is 100x (Gnerre et al. 2011). Recently, Mohr and colleagues (Mohr et al. 2017) presented a highly continuous assembly of the endangered Hawaiian Monk seal (∼2.4Gb total genome assembly length) using a combination of 10x Genomics Chromium and Bionano Genomics optical mapping. Interestingly, their 10x Genomics Chromium assembly showed similar N50 statistics to those reported here (scaffold N50 22.23Mb), showing that 10x Genomics Chromium technology alone enables the generation of high-quality mammalian genome assemblies.

A limitation of 10x Genomics Chromium technology is the requirement of fresh tissue samples for the isolation of HMW DNA. This can be difficult or impossible to obtain from some endangered species. Fortunately, small amounts of mammalian blood yield sufficient amounts of HMW DNA when properly stored and can be sampled without causing harm to the animal. Additionally, DNA extraction kits such as the Qiagen MagAttract kit can extract sufficient amounts of HMW DNA from as little as 200μl. For museum samples, or tissues stored for extended periods of time, reference-based mapping might be the only option to extract long-range genomic information. However, for extant endangered species, especially those with individuals in captivity, 10x Genomics Chromium offers a cost-effective approach to sequence genomes. For species with genome sizes <1Gb and between ∼3Gb and 5.8Gb special data processing will need to be applied (see https://support.10xgenomics.com/de-novo-assembly/sample-prep/doc/technical-note-supernova-guidance). In addition, the amplification primers for the 10x Chromium library preparation are designed for GC contents similar to human (∼41%), implying that the method might not work as well for genomes that strongly divert from this GC content (e.g. for some invertebrates).

### Cost effectiveness

Sequencing costs are steadily dropping. At the time the sequencing for this project was carried out a lane on the Illumina HiSeqX cost approximately $1,500 - $2,000 and a 10x Genomics library ranged from $450 to $1000, thus allowing the generation of high quality *de novo* genomes for less than $3,000 total. Even more so, independent of sequencing lane costs, this method only requires a single library to be sequenced to an average coverage of 25 - 75x, unlike other methods which require multiple libraries at higher coverage. As we have shown here, continuous assemblies can be generated from as little as 25x. Furthermore, computational resources required to assemble the genome are very low. The current version of Supernova 1.2 only requires a minimum of 16 CPU cores and 244Gb of memory (for a human genome at 56x coverage; http://www.10xgenomics.com/), and the assembly can be carried out in only few days (depending on the number of available CPU cores). This is about a reduction of five times the memory requirement compared to the first version of Supernova. Even more so, Supernova does not require parameter input or tuning, thus allowing even novices to easily assemble 10x Genomics Chromium based genomes.

### Applications in conservation

Traditionally, conservation biologists have obtained a great deal of genetic information from a few microsatellite markers and/or nuclear and mitochondrial loci. The analysis of microsatellite markers can provide a snapshot into contemporary population structure, but this method risks providing incomplete information on selection and migration and it is not a reliable way to identify individuals due to the stochastic behavior of marker amplification (Taberlet & Luikart (1999), reviewed in Morin et al. (2004)). Moreover, microsatellites can be difficult to successfully design and develop, which can quickly increase costs for species that have little to no genetic information available. The ability to rapidly and cost-effectively generate full genomes will allow conservation biologists to bridge this gap and harvest crucial fine-scale population information for population parameters such as inbreeding (e.g. Vieira et al. (2013)), load of deleterious mutations (e.g. Robinson et al. (2016)), gene flow (e.g. Pazmiño et al. (2017)) and population structure (e.g. Hampton et al. (2004)). Once a reference genome has been assembled, optional (low coverage) re- sequencing data from several individuals allows for the typing of genome-wide information such as single-nucleotide polymorphisms (SNPs), potentially neutral microsatellite loci, and other genomic regions of interest. These data can then be used to investigate the abovementioned population parameters, but also further yield insights into adaptive genetic variation and perhaps the adaptive potential of different populations or species. Furthermore, genome-wide SNP or mapping data can help us to reconstruct recent and ancient population histories, using methods such as PSMC (Li & Durbin 2011), MSMC (Schiffels & Durbin 2014), and Stairway plots (Liu & Fu 2015). These questions have gone largely unanswered for many species, but warrant investigation so we can better understand how humans have affected species contemporary distributions and what their suitable habitats might have looked like.

### Heterozygosity within African Wild dog individuals

A high number of heterozygous sites were shared between all three individuals in this study, with Sister 1 and Sister 2 sharing more heterozygous sites than either with Eureka. Each of the individuals further shows a high number of singletons (heterozygous sites only found in one individual). Even when compared to the two low coverage genomes from Campana et al. (2016) we find a high number of shared sites. As expected, we see a much higher rate of singletons in these two individuals. Due to the low coverage (5.7 - 5.8x average coverage) we predict a higher proportion of the called heterozygous sites to be false positives due to sequencing errors. Heterozygosity per site estimates indicate a high within individual diversity. Estimates ranged from 0.0007 - 0.001 for Eureka to 0.0009 - 0.0012 for Sister 2. Intriguingly, other threatened mammals, such as the Iberian lynx *(Lynx pardinus*), the cheetah (*Acinonyx jubatus*) or the island fox (*Urocyon littoralis*) show nearly 10 fold lower heterozygosity (0.0001 (Abascal et al. 2016), 0.0002 (Dobrynin et al. 2015) and 0.000014 - 0.0004 (Robinson et al. 2016), respectively). The high within-individual heterozygosity could be a result of their social structure, as only unrelated individuals come together to form new packs through dispersal. This could be very good news for the survival of these species if external pressures (such as hunting, habitat fragmentation, etc.) can be reduced.

The Supernova software reports distance between heterozygous site estimates (see Supplementary Table 1). Interestingly, those estimates were much lower than the ones obtained based on the genotype calls produced with ANGSD. While Supernova estimated this distance to be 2.6kb in Sister 1, 3.1kb in Sister 2 and 7.1kb in Eureka, the ANGSD based estimates range from 850bp - 1.2kb for Sister 1, 814bp - 1.1kb for Sister 2 and 999bp - 1.5kb for Eureka, depending on the posterior cutoff used. Supernova calculates the distance between heterozygous sites as part of the assembly process. However, when the fasta consensus sequence is called part of the variation can get flattened (see Weisenfeld et al. (2017)). This can happen in regions between mega bubbles, which are nominally homozygous, but could actually have some variation that cannot be phased by Supernova. This could explain the lower heterozygosity values. However, we should point out that heterozygosity values obtained using genotype calls in ANGSD could also be biased, as they are based on the nominal and not the effective coverage. The nominal coverage is the total number of reads that cover a site in the assembly, whereas for the effective coverage only reads from different barcodes are included in the estimation. If individual barcoded regions amplified with different efficiency during the library preparation step, then heterozygosity estimates could be unreliable. However, this should not strongly affect genome-wide heterozygosity estimates, as we expect this issue to be rare.

## Conclusion

We find that the 10x Genomics Chromium system can be used to assemble highly continuous and accurate mammalian genome assemblies for less than $3,000 US dollars per genome (sequenced 2016 and 2017). The method can be easily applied to species of conservation concern for which genomic methods could greatly benefit their management and monitoring programs. For the African wild dog, these genomes will facilitate more reliable and cost-effective conservation efforts through the use of re-sequencing and SNP-typing methods. Compared to other species of conservation concern, the African wild dog has a relatively high heterozygosity. More studies are required to understand how both the social biology and recent precipitous population declines have impacted the population genomic structure of African wild dogs, and how management might use this information for the benefit and longevity of the species.

## Acknowledgements

We thank Mary Agnew, Cheryl Asa, Luis Padilla, and Wessly Warren for assistance in obtaining the Eureka sample. Tyler Linderoth, Thorfinn Korneliussen, and Ke Bi for help with the different heterozygosity calculations and interpretations. Deanna Church from 10x Genomics for discussion on how SuperNova performs the heterozygous site calling.

## Supplemental Info

**Supplementary Figure S1.**
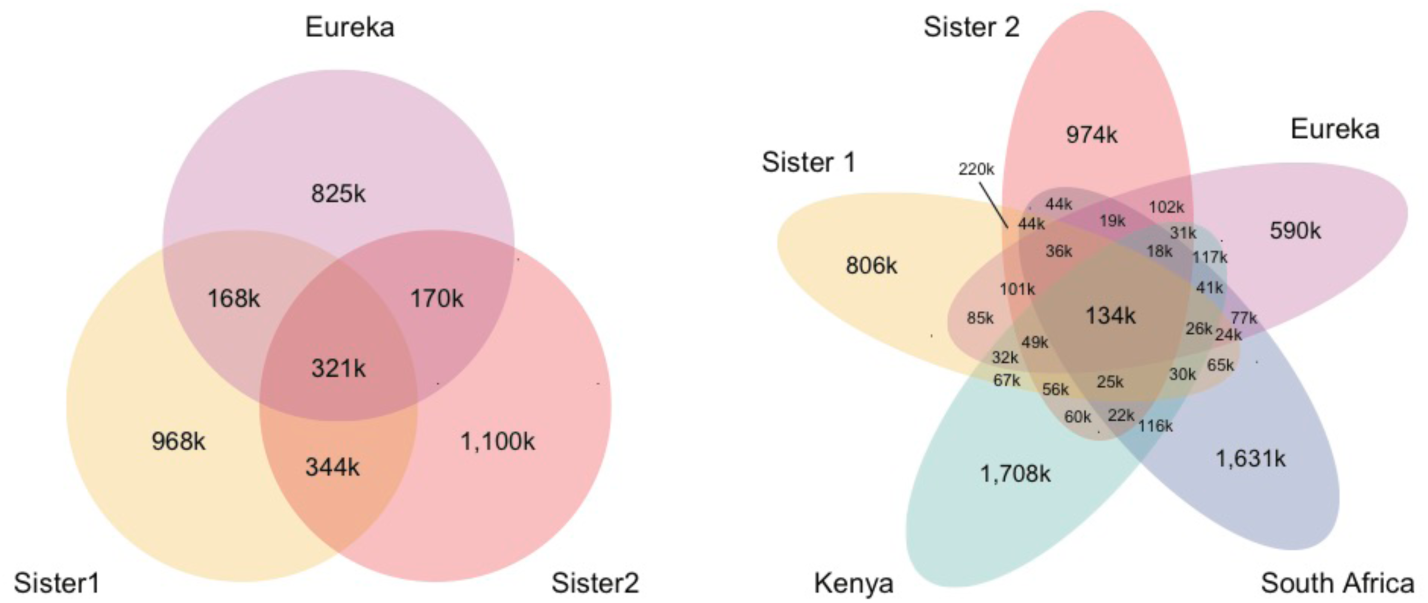
Comparison of heterozygous sites between individuals. A) Shared heterozygous sites between the three *de novo* assemblies. Many heterozygous sites are shared between all individuals, and more heterozygous sites are shared between the two sisters than between each sister and Eureka. Same plot as Fig. 1B in the main manuscript) Shared heterozygous sites between the three *de novo* assemblies and the two low-coverage reference-based genomes (Kenya and South Africa) from Campana et al. 2016. Both Kenya and South Africa show a very high number of singletons, which is likely caused by the low coverage and the resulting false-positive heterozygous sites (caused by sequencing errors). We see that a high amount of heterozygous sites are shared between all individuals, and that Sister 1 and Sister 2 share more heterozygous sites than any other pairwise comparison.

**Supplementary Figure S2.**
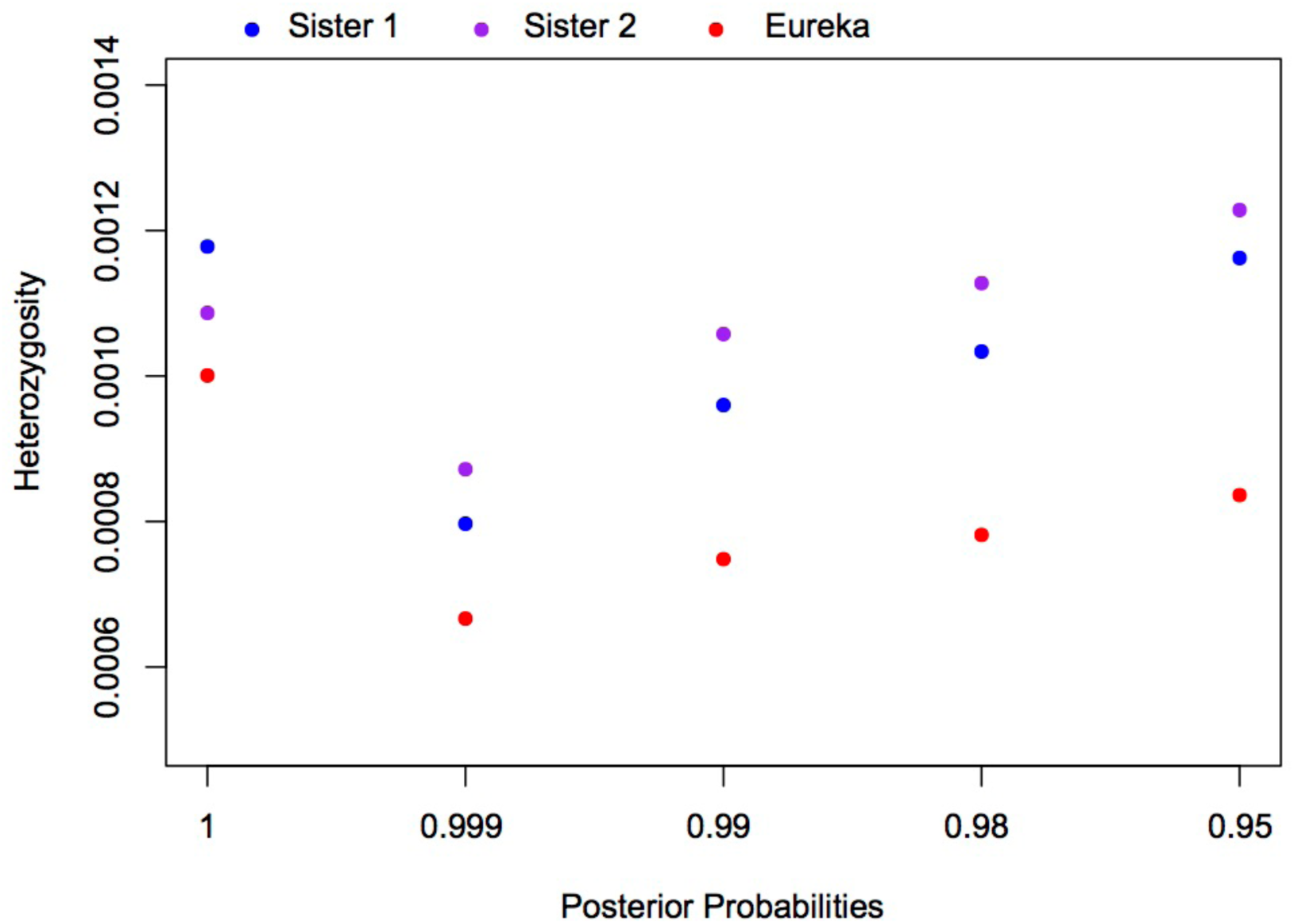
Comparison of heterozygosity estimates using different posterior probability cutoffs for all three assemblies. We used an average coverage of 20x for the heterozygosity estimations.

**Supplementary Table S1.** Assembly statistics as calculated by Supernova.

| Statistic | Sister 1 | Sister 2 | Eureka |
| --- | --- | --- | --- |
| Reads | 1200 M | 801.56 M | 427.6 M |
| Mean read length | 138 bp | 139 bp | 138 bp |
| Read two Q30 | 71.56 % | 87.01 % | 80.86 % |
| Median insert | 0.38 Kb | 0.31 Kb | 0.34 Kb |
| Proper pairs | 86.04 % | 89.44 % | 86.81 % |
| Molecule length | 19.91 Kb | 77.03 Kb | 52.0 Kb |
| Heterozygosity distance | 2.61 Kb | 3.11 Kb | 7.14 Kb |
| Number of unbarcoded reads | 5.79 % | 4.77 % | 5.17 % |
| N50 reads per barcode | 972 | 678 | 354 |
| **Duplicates** | 23.46 % | 21.7 % | 3.28 % |
| **Phased** | 39.07 % | 40.1 % | 52.54 % |
| **Scaffolds >= 10kb** | 1.12 K | 1.2 K | 1.56 K |
| **N50 edge size** | 5.96 Kb | 10.88 Kb | 9.36 Kb |
| **N50 contig size** | 61.34 Kb | 83.47 Kb | 50.15 Kb |
| **N50 phase block size** | 0.12 Mb | 2.02 Mb | 0.31 Mb |
| **N50 scaffold size** | 7.91 Mb | 21.34 Mb | 15.31 Mb |
| **N60 scaffold size** | 6.2 Mb | 17.04 Mb | 11.49 Mb |
| **Assembly size (scaffolds >=10kb)** | 2.27 Gb | 2.26 Gb | 2.20 Gb |

**Supplementary Table S2.** Repeat statistics. *De novo* and homology based repeat annotations as reported by RepeatMasker and RepeatModeler. Families of repeats included here are long interspersed nuclear elements (LINEs), short interspersed nuclear elements (SINEs), long tandem repeats (LTR), DNA repeats (DNA), unclassified (unknown) repeat families, small RNA repeats (SmRNA), and others (consisting of small, but classified repeat groups). The total is the total percentage of base pairs made up of repeats in each genome, respectively.

| **Assembly** | **LINE** | **SINE** | **LTR** | **DNA** | **Unclassified** | **SmRNA** | **Others** | **Total (%)** |
| --- | --- | --- | --- | --- | --- | --- | --- | --- |
| Sister 1 | 847,645 | 1,472,018 | 297,851 | 315,865 | 11,125 | 1,080,087 | 1,054,219 | 40.19 |
| Sister 2 | 857,045 | 1,490,757 | 301,853 | 319,940 | 10,103 | 1,095,228 | 1,050,189 | 41.42 |
| Eureka | 855,470 | 1,479,441 | 299,492 | 317,939 | 12,936 | 1,086,289 | 1,025,874 | 38.65 |
| canFam3.1 | 857,579 | 1,503,465 | 302,932 | 321,141 | 14,466 | 1,110,467 | 1,038,344 | 42.13 |

**Supplementary Table S3.**
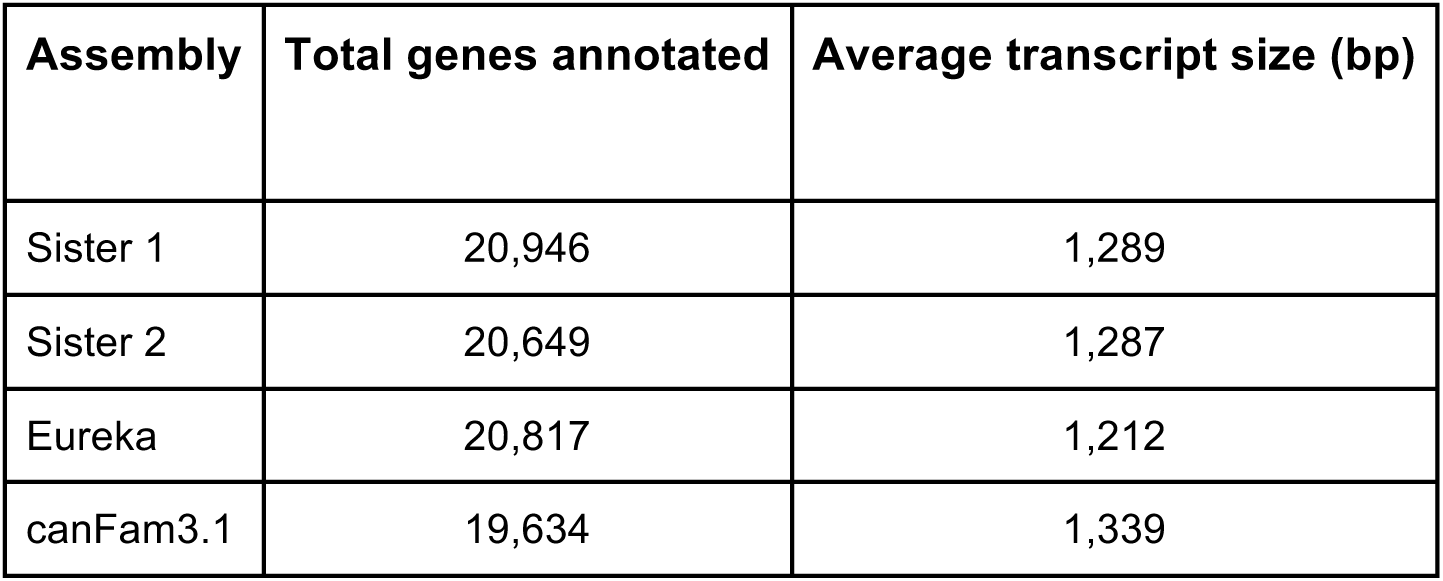
Gene Annotation. Total number and average gene transcript sizes as reported by Maker3.

| **Assembly** | **Total genes annotated** | **Average transcript size (bp)** |
| --- | --- | --- |
| Sister 1 | 20,946 | 1,289 |
| Sister 2 | 20,649 | 1,287 |
| Eureka | 20,817 | 1,212 |
| canFam3.1 | 19,634 | 1,339 |

**Supplementary Table S4.** Heterozygosity estimates. The total number of sites, the total number of heterozygous sites, the calculated heterozygosity and the length in bp between heterozygous sites is provided for all three genomes for different posterior probability cutoffs.

| Genome assembly | Posterior cutoff | Total number of Sites | Total number of heterozygous sites | Heterozygosity | Length (bp between heterozygous sites) |
| --- | --- | --- | --- | --- | --- |
| **Sister 1** | 1 | 940,480,720 | 1,107,829 | 0.0012 | 849 |
|  | 0.999 | 1,863,729,959 | 1,485,163 | 0.0008 | 1,255 |
|  | 0.99 | 1,876,594,049 | 1,801,440 | 0.0010 | 1,042 |
|  | 0.98 | 1,879,378,076 | 1,942,525 | 0.0010 | 967 |
|  | 0.95 | 1,883,014,319 | 2,188,413 | 0.0012 | 860 |
| **Sister 2** | 1 | 1,063,807,193 | 1,156,011 | 0.0011 | 920 |
|  | 0.999 | 1,820,298,841 | 1,586,917 | 0.0009 | 1147 |
|  | 0.99 | 1,829,356,947 | 1,934,764 | 0.0011 | 946 |
|  | 0.98 | 1,831,414,501 | 2,065,053 | 0.0011 | 887 |
|  | 0.95 | 1,833,781,723 | 2,252,213 | 0.0012 | 814 |
| **Eureka** | 1 | 1,123,979,892 | 1,124,799 | 0.0010 | 999 |
|  | 0.999 | 1,972,583,001 | 1,314,557 | 0.0007 | 1,501 |
|  | 0.99 | 1,983,203,305 | 1,483,893 | 0.0007 | 1,336 |
|  | 0.98 | 1,985,824,623 | 1,551,950 | 0.0008 | 1,280 |
|  | 0.95 | 1,989,156,093 | 1,663,356 | 0.0008 | 1,196 |

